# Antibody-Free Immunopeptide Nano-Conjugates for Brain-Targeted Drug Delivery in Glioblastoma Multiforme

**DOI:** 10.1101/2025.03.07.641755

**Authors:** Saurabh Sharma, David Lee, Surjendu Maity, Prabhjeet Singh, Jay Chadokiya, Neda Mohaghegh, Alireza Hassani, Hanjun Kim, Ankit Gangarade, Julia Y. Ljubimova, Amanda Kirane, Eggehard Holler

**Affiliations:** Department of Surgery, Division of Surgical Oncology, Stanford School of Medicine, Stanford University Medical Center, CA 94305, USA; Terasaki Institute for Biomedical Innovation, Los Angeles, CA 90064, USA; Department of Orthopedic Surgery, Duke University School of Medicine, Duke University, Durham, NC 27710, USA

**Author notes:** Dr. Amanda Kirane, MD, PhD, FACS, FSSO, Assistant Professor of Surgery, Department of Surgery, Director, Cutaneous Surgical Oncology, Stanford School of Medicine, CA-USA-94305 Email-, Eggehard Holler, PhD, Professor, Terasaki Institute of Biomedical Innovation, Los Angeles, 90024, USA. These authors have contributed equally to this work (Amanda Kirane, Eggehard Holler).

**Keywords:** Nanoconjugate, Immunotherapy, 3D Tumor-BBB Model, Biodistribution

## Abstract

Glioblastoma Multiforme (GBM) represents a significant clinical challenge amongst central nervous system (CNS) tumors, with a dismal mean survival rate of less than 8 months, a statistic that has remained largely unchanged for decades (National Brain Society, 2022). The specialized intricate anatomical features of the brain, notably the blood-brain barrier (BBB), pose significant challenges to effective therapeutic interventions, limiting the potential reach of modern advancements in immunotherapy to impact these types of tumors. This study introduces an innovative, actively targeted immunotherapeutic nanoconjugate (P12/AP-2/NCs) designed to serve as an immunotherapeutic agent capable of traversing the BBB via LRP-1 receptor-mediated transcytosis. P12/AP-2/NCs exert its immune-modulating effects by inhibiting the PD-1/PD-L1 axis through a small-size PD-L1/ PD-L2 antagonist peptide Aurigene NP-12 (P12). P12/AP-2/NCs are synthesized from completely biodegradable, functionalized high molecular weight β-poly(L-malic acid) (PMLA) polymer, conjugated with P12 and Angiopep-2 (AP2) to yield P12/AP-2/NCs. Evaluating nanoconjugates for BBB permeability and 3-D tumor model efficacy using an in vitro BBB-Transwell spheroid based model demonstrating successful crossing of the BBB and internalization in brain 3D tumor environments. In addition, the nanoconjugate mediated T cell’s cytotoxicity on 3D tumor region death in a U87 GBM 3-D spheroid model. AP2/P12/NCs is selectively inhibited in PD1/PDL1 interaction on T cells and tumor site, increasing inflammatory cytokine secretion and T cell proliferation. In an in-vivo murine brain environment, rhodamine fluorophore-labeled AP2/P12/NCs displayed significantly increased accumulation in the brain during 2-6 h time intervals post-injection with a prolonged bioavailability over unconjugated peptides. AP2/P12/NCs demonstrated a safety profile at both low and high doses based on major organ histopathology evaluations. Our findings introduce a novel, programmable nanoconjugate platform capable of penetrating the BBB for directed delivery of small peptides and significant immune environment modulation without utilizing antibodies, offering promise for treating challenging brain diseases like glioblastoma multiforme and beyond.

## 1. Introduction

Glioblastoma multiforme (GBM) is considered the most aggressive and lethal form of primary brain tumor. A grim prognosis, high mortality, and increased recurrence rates are characteristics of GBM. Communication within the tumor-immune microenvironment (TiME) is the main determinant factor for its aggressiveness^1^. GBM cells recruit immunomodulatory cells such as macrophages, microglia, neutrophils, mast cells, T cells, lymphatic vessels, and the extracellular matrix to promote cancer cell invasiveness and dissemination by stimulating angiogenesis, suppressing immune cell functions, and altering cancer cell metabolism^2^. Furthermore, multiple biological barriers in GBM, especially the blood-brain barrier (BBB) greatly hinder the effective delivery of therapeutic agents. The BBB is formed by specialized brain capillary endothelial cells, along with pericytes and astrocytes, which work together to protect the brain from harmful substances and maintain its stability^3^. Because of this, only a limited number of drugs can penetrate the BBB. Due to these variables, the median survival time of GBM patients is only 12–15 months after their diagnosis^4^. Therefore, it’s crucial to devise innovative, multifunctional drug delivery systems that can effectively navigate both the BBB and BBTB to precisely target glioma cells and enhance the effectiveness of the treatments.

Immune checkpoint blockade (ICB) therapy has emerged as a groundbreaking treatment option for combating tumors. However, the effectiveness of ICB therapy, particularly PD-L1 antibody-based approaches in treating brain tumors, has been limited. This is largely due to the blood-brain barrier (BBB), which restricts the transit of ICB antibodies into the brain tissue. Additionally, while the most commonly used anti-PD-1, CTLA-4 therapies can stimulate an immune response against certain tumors, they can also lead to serious immune-related adverse events (irAEs) in the peripheral regions^5^. These irAEs can be severe, affecting up to 43% of patients treated with CTLA4 antibodies and around 20% with PD-1/PD-L1 antibodies^6,7^. For several major cancer types, including colorectal cancer, anti-PD-1/PD-L1 treatment demonstrates minimal efficacy^8^. The likelihood of experiencing irAEs increases with higher doses, as the toxicity correlates with the amount of antibodies administered^9^. The long half-life of these antibodies, which can exceed 15-20 days, and their prolonged target occupancy—often lasting for months—contribute to the severe irAEs observed in clinical settings^10,11^. To address these challenges, researchers are exploring small peptides as a promising alternative for ICB therapy. Peptides offer several advantages, including similarity to natural proteins, a high level of specificity toward their targets, and reduced toxicity compared to traditional compounds. In this research, we have used Aurigene NP-12 (P-12), a peptide designed to act as an immune checkpoint antagonist of PD-L1. Composed of 29 amino acids, P-12 is a decoy for PD-1 derived from specific regions of the human PD-1 receptor^12^. P-12 showed moderate clearance and a relatively low volume of distribution. Peak plasma concentrations were attained between 0.2 and 0.4 hours following subcutaneous administration. The absolute bioavailability in mice was shown to be 77%. The B16F10 mouse melanoma spontaneous lung metastasis model received P12 subcutaneously at a dosage of 3 mg/kg/day for 14 days. P-12 treatment led to a 66% decrease in metastatic nodules relative to vehicle control. One significant characteristic of P-12 is its relatively short half-life of approximately 0.22 h, which may lead to increased frequency of administration but could also translate to fewer side effects, such as irAEs, associated with conventional ICB therapies^12^.

Given that glioblastoma (GBM) cells express PD-L1^13^, employing P-12 in treatment strategies utilizing nanoconjugate technology could represent a logical and effective approach to target this aggressive brain tumor, which is still an unknown territory for research. So, this will be the first reported scenario where we will target P-12 peptide specifically for GBM. Another hurdle hindering the effective delivery of NP-12 for GBM is the BBB and the BBTB. To address this challenge, our team has synthesized a biodegradable nontoxic β-poly(L-malic acid) (PMLA), which possesses polymeric pendent carboxylic acid groups to bind the BBB chemically crossing peptides P-12 and Angiopep-2 (AP2-LRP-1 binding ligand)^14,15^. Certain molecules can cross the blood-brain barrier (BBB) through specific transport mechanisms. One such example is the low-density lipoprotein receptor pathway (LRP-1), which facilitates the movement of low-density lipoproteins into and out of the brain. Accumulating evidence reveals that LRP1 expressed on the abluminal site is the major mechanism involved in the brain-to-blood efflux of soluble β-amyloid peptides with the help of the apolipoprotein family^16^. When ligands attach to LRP-1 in blood vessel cells, they are internalized and transported into the brain tissue.

A synthetic peptide ligand, Angiopep-2 (AP2; TFFYGGSRGKRNNFKTEEY), has been developed, which saturates the LRP-1 receptors by inhibiting other LRP-1 ligands^15^. This confirms transcytosis of AP2 across the BBB. In GBM, there has been a clear clinical correlation between LRP-1 and its role in tumor progression^17^. Based on this data, BBB-GBM models have observed that LRP-1 showed an upregulation in the vasculature near GBM spheroids. Studies by Straehla et al. showed that incorporating Angiopep-2 (AP2) peptide moieties on the surface of NPs leads to increased BBB permeability near GBM tumors throughLRP1-mediated transcytosis^18^. In addition, a tri-leucine (LLL) chain for hydrophilicity and endosomal escape unit was attached, and a fluorescent marker Rhodamine (Rh) was attached to the PMLA backbone. We consider the PMLA-tri leucine group amphiphilic, which contains multiple carboxylic groups for vector-receptor attachment and can be biodegraded by lipases and peptidases, thereby escaping deposit toxicity in vivo.

In this research, we explore the efficacy of Angiopep-2 and P-12 PMLA-based nanoconjugates (AP-2/P-12/NCs) for the targeted delivery of P-12 peptides for treating GBM. Our findings indicate that this polymer nanoconjugate not only crosses the blood-brain barrier through receptor-mediated uptake but also disrupts PD-1/PD-L1-mediated immune evasion with the help of the P-12 peptide. This dual action fosters a more robust anti-tumor immune response within the brain tumor immune-microenvironment. We utilized a human-derived BBB transwell model with GBM spheroids to assess the efficacy of the developed nanoconjugates, complemented by comprehensive T-cell activation and cytokine profiling. Furthermore, our in vivo and ex vivo biodistribution and tissues toxicity studies in C57BL/6 mice provide additional evidence of the potential of these nanoconjugates as a transformative approach in the GBM immunotherapy landscape. The proposed AP-2/P-12/NCs demonstrate the potential for enhancing bioavailability while minimizing systemic toxicity, paving the way for more effective immunotherapy strategies for brain cancer.

## 2. Results

### 2.1 Synthesis and Characterization of AP-2/P-12 functionalized Nanoconjugates (AP-2/P-12 NCs)

The AP-2/P-12 immunotherapeutic NCs were synthesized using polymalic acid extracted from sp. *Physarum polycephalum* and purified to yield pure polymer with an average molecular weight of 50-100 kDa^15^ as we published earlier ^15, 23–29^. Due to its biocompatibility and ease in tailorable nature, the PMLA possessing free COOH groups were utilized as a polymer backbone for our immune-nanoconjugate (Fig. 2A). The purified biosynthesized PMLA polymer was reacted with DCC/NHS to activate the free COOH sites of the polymer, followed by reaction with tri leucine (LLL), P-12 peptide (PD-L1 blocker) or FITC-P-12 peptide, and mercaptoethylamine (MEA) to obtain pre-conjugate P-12/thiolated PMLA. The pre-conjugate with the free thiol (SH) group was reacted with pre-functionalized maleimide-PEG-AP-2 peptide (LRP1 peptide; **(1)**) and PDP **(2)** to obtain AP-2/P-12 NCs (Fig. 2A). The fluorescently labeled AP-2/P-12 NCs were prepared by reacting Rhodamine Red™-C2-maleimide with AP-2/P-12 NCs to obtain AP-2/P-12/RhB NCs. AP-2 and P-12 peptide conjugation efficiency was determined during the synthesis process using size exclusion high-pressure liquid chromatography (SEC-HPLC). The resulting data showed that the average molecular weight of PMLA and AP-2/P-12 NCs was 67 KDa and 145 KDa, respectively (Fig. 2B). The conjugation of AP-2 and P-12 peptide was confirmed using 1H NMR, followed by the characterization using Zetasizer Nano ZS90 (Malvern Instruments, Malvern, UK), indicating an average particle size of less than 100 nm, and polydispersity index of < 0.3 (Supplementary Table 1).

**Figure 1.**
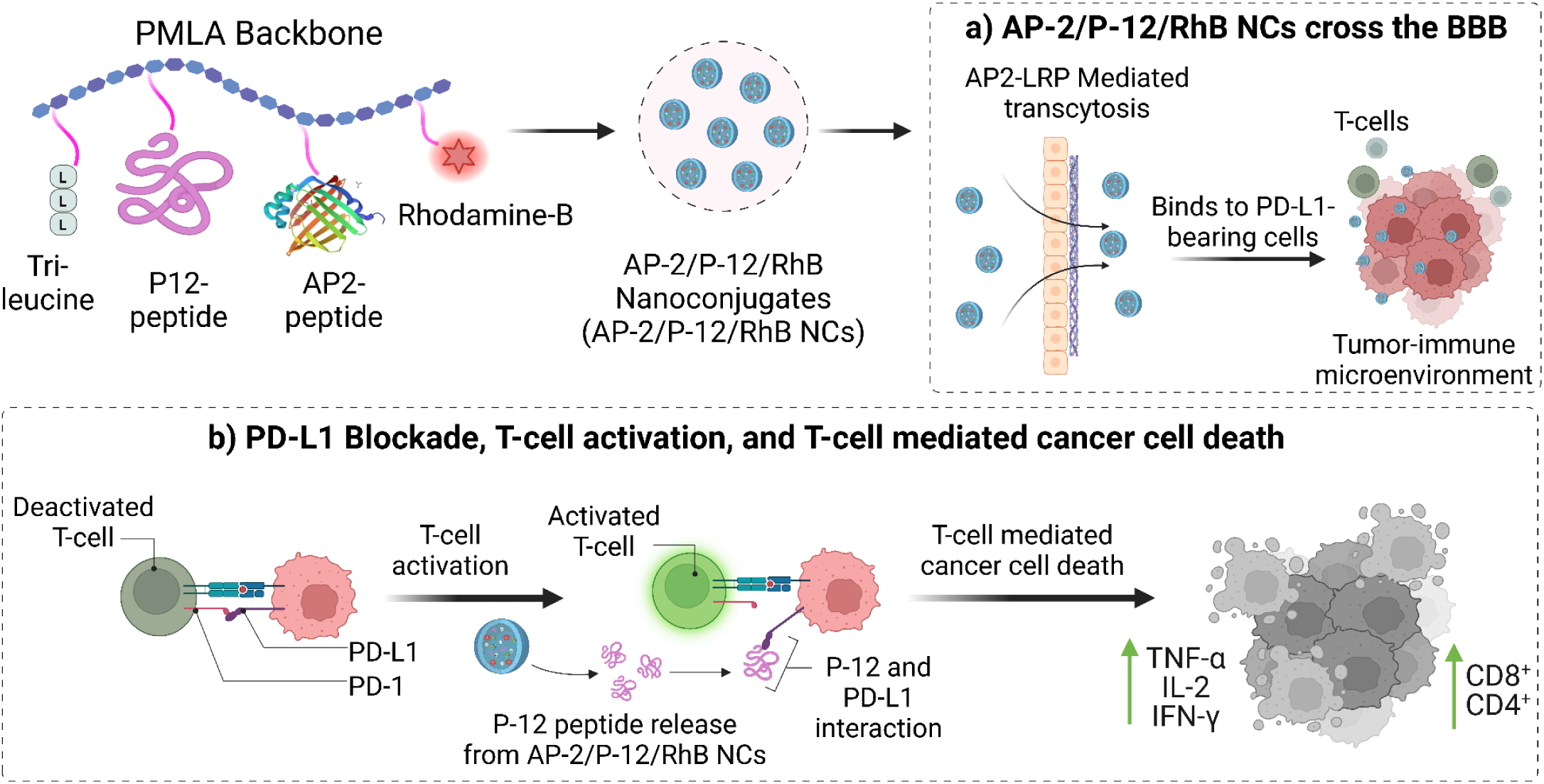
The introductory mechanism of action for AP-2/P-12/RhB nanoconjugates (NCs) involves a multi-step process that enhances their therapeutic efficacy against brain tumors. In a) the AP-2 in the nanoconjugates utilize LRP1 receptor-mediated transcytosis to cross the BBB and BBTB. This process allows the NCs to reach the brain tumor site. Once at the tumor site, b) the P-12 blocks the PD-1/PD-L1 immune checkpoint interaction. This blockade occurs between T cells and PDL1-positive brain tumor cells, preventing the tumor cells from evading immune surveillance. This disruption of the PD-1/PD-L1 axis leads to enhanced T cell activity, characterized by: i) Increased T cell (CD4+/CD8+ Cells) proliferation and ii) Upregulation of major proinflammatory cytokine expression such as TNFα, IL-2, IFN-γ. The activated T cells exhibit heightened cytotoxic activity against the brain tumor cells, promoting tumor cell death and potentially reducing tumor burden.

**Figure 2.**
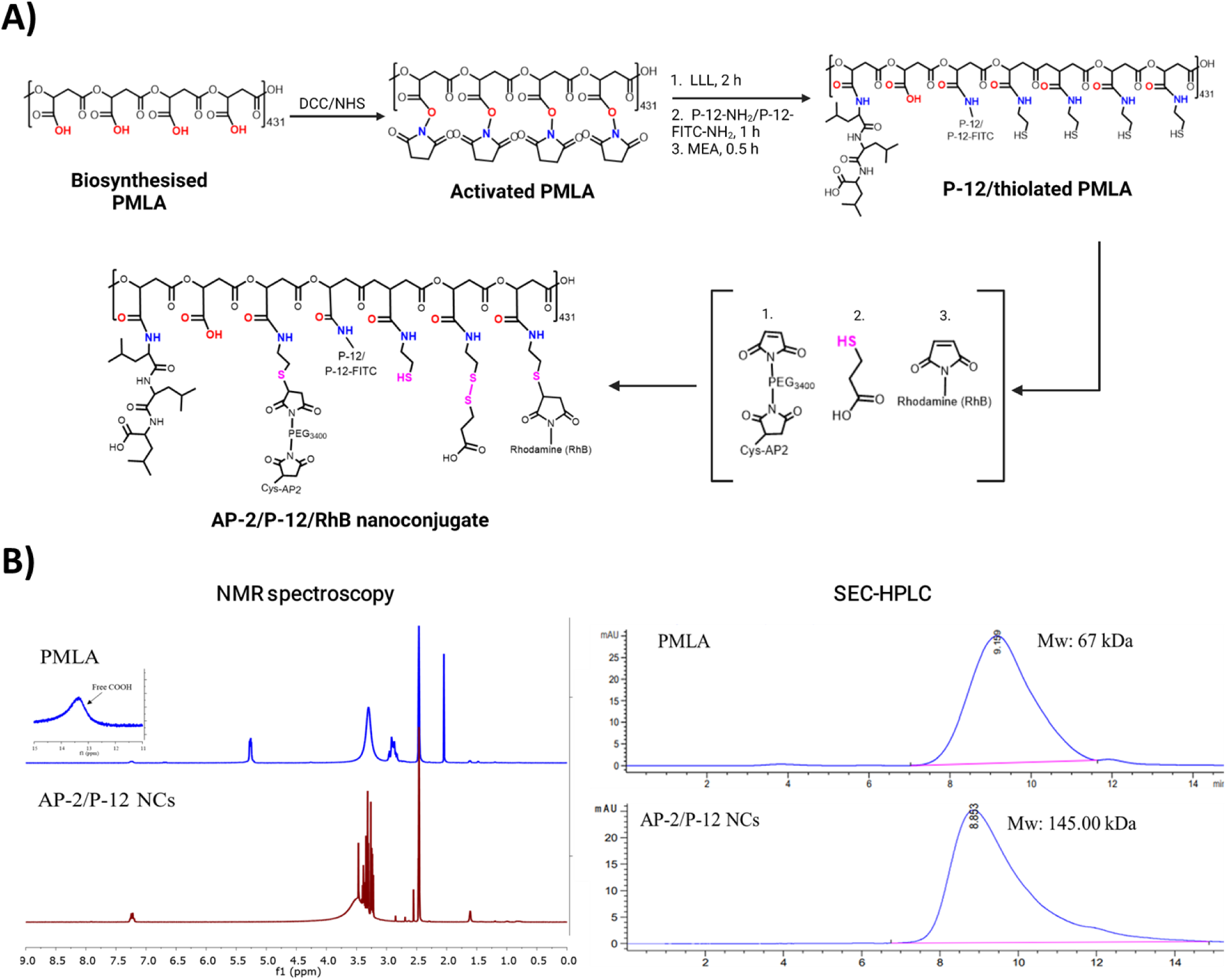
Synthesis of AP-2/P-12/RhB NCs: A) Synthesis of AP2-2/P-12/Rhb NCs from PMLA which is a multistep process involving the activation of PMLA obtained from a biosynthetic source, which will render the PMLA backbone to conjugate the AP-2, P-12, and RhB moieties successfully. B) Characterization by using 1H NMR to confirm successful conjugation of functional moieties to the PMLA backbone and SEC-HPLC for determining the total molecular weight distribution of the synthesized PMLA and AP-2/P-12 NCs.

### 2.2 AP-2/P-12 NCs crosses the human-derived BBB transwell model with GBM tumor spheroids model in vitro

AP-2/P-12 NCs penetration efficiency was determined using BBB GBM spheroid model. Wherein human brain endothelial cells (HBMEC) coated transwell with U87MG-based humanoid 3D GBM spheroids were prepared to mimic the BBB properties (Fig. 3A). The developed HBMEC BBB spheroid model was validated using immunocytochemistry (ICC) for their characteristic protein marker such as zona occludens-1 (ZO-1) (red color) for protein located on a cytoplasmic membrane surface of intercellular tight junctions, VE-cadherin (green color) for the protein localized at the intercellular boundaries of endothelial cells, GLUT-1 (red color) protein for facilitative glucose transporter, and CD31 (green color) for platelet endothelial cell adhesion molecule-1 (PECAM-1) protein (Supplementary Fig. S2). The ability of the Rhodamine conjugated AP-2/P-12 NCs (AP-2/P-12/RhB NCs) to cross the BBB was evaluated by using intracellular uptake assay in BBB layer using the confocal microscopy (Fig. 3B). Due to the presence of AP-2 peptide in AP-2/P-12/RhB NCs, it underwent LRP1 receptor-mediated transcytosis, exhibiting better cellular uptake potential as depicted with improved rhodamine signal intensity compared to free rhodamine (control) in HBMEC (Fig. 3B). AP-2/P-12/RhB NCs penetration potential was evaluated in the U87MG-based 3D-GBM spheroid model (Fig. 3C). The treatment with AP-2/P-12/RhB NCs containing rhodamine exhibited higher signal intensity in both the core (∼7 folds) and periphery (∼12 folds) of the U87 tumor spheroid compared to the free rhodamine-treated group (Fig. 3 C & D).

**Figure 3.**
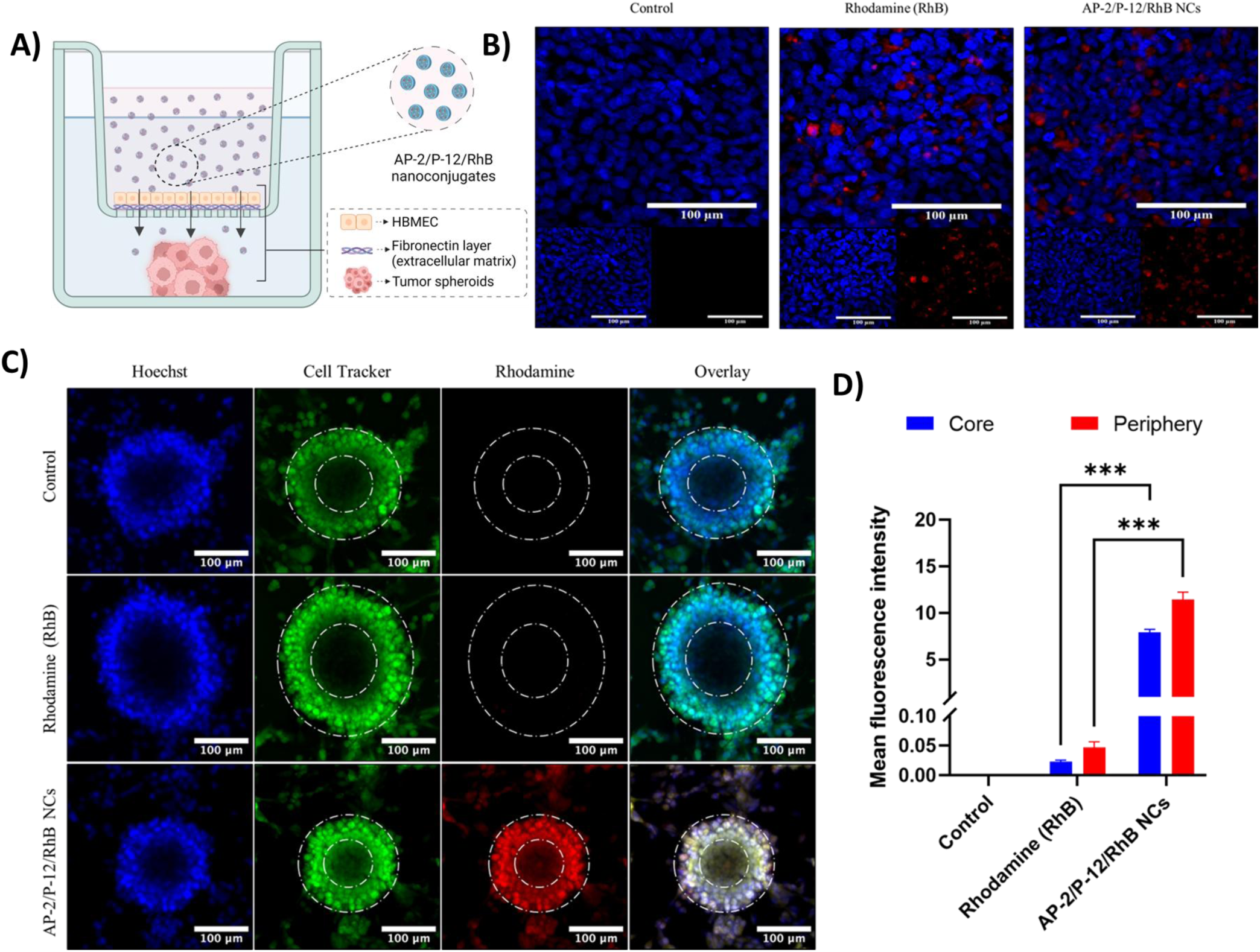
A) Schematic representation of a BBB-transwell model demonstrating nanoparticle receptor-mediated transcytosis and PDL1 receptor-mediated interaction with GBM spheroids wherein we have incorporated human brain microvascular endothelial cells (HBMECs) on a fibronectin layer to mimic the blood-brain barrier (BBB). The model illustrates the successful crossing of AP-2/P-12/RhB NCs through the BBB and their subsequent interaction with glioblastoma multiforme (GBM) tumor spheroids in a separate compartment. B) Cellular uptake of nanoconjugates in a BBB transwell model: Confocal laser scanning microscopy (CLSM) analysis of AP-2/P-12/RhB NCs uptake by HBMECs (seeded as a monolayer on fibronectin-coated transwells) mimicking the BBB and incubated with 0.1 mg/mL of AP-2/P-12/RhB NCs for 2 h. C) Nanoconjugate penetration in 3D GBM spheroid model carried out by CLSM visualization of AP-2/P-12/RhB NCs (100 ug/mL) uptake and distribution within 3D U87 glioblastoma spheroids after 4 h incubation at 37°C. D) Quantitative analysis of nanoconjugate uptake carried out by measuring the mean fluorescence intensity from CLSM images, representing the relative uptake and accumulation of AP-2/P-12/RhB NCs in both the BBB model and GBM spheroids. Data are presented as mean ± standard error of the mean (SEM). Statistical significance was determined using one-way ANOVA followed by post-hoc tests, with ** p < 0.01, *** p < 0.001, and **** p < 0.0001 indicating significance levels between treatment groups.

### 2.3 AP-12/P-12 NCs induce the proliferation and activation of proinflammatory T cells

The proliferation potential of AP-2/P-12 NCs on the immune cells in highly immunosuppressive PDL1/PDL2-rich environments was evaluated by using CD3/CD28 activated T-cells. The isolated T-cells were stimulated with anti-CD3/CD28 antibodies and then treated with CFSE (Fig. 4A). After that, the activated T-cells treated with PDL1/L2 protein and free P-12-pep resulted in increased T-cell proliferation in both CD4/CD8+ T-cells. Further, the treatment with PDL1/L2 protein and AP-2/P-12 NCs exhibited an additional improvement in CD8+/CD4 (p<0.001) proliferation potential with minimal impact FOXP3+ expressing regulatory T-cells (Treg) (Fig. 4B). Simultaneously, the culture media was evaluated for various pro-inflammatory cytokines including IL-2, IFN-γ, and TNF-α. Wherein the treatment with free P-12-pep indicated improvement in the IL-2, IFN-γ, and TNF-α expression compared to control and αPD-1 antibody; but upon treatment with AP-2/P-12 NCs resulted in further improvement in the pro-inflammatory cytokine expression compared to free P-12-pep counterpart (Fig. 4C). Overall, the improved expression of T-cells, and pro-inflammatory cytokines indicated an improved checkpoint blocking capability in between PD1-PDL1/L2 complex formation. They could be utilized as a better strategy for targeting PDL1/L2-expressing tumors.

**Figure 4:**
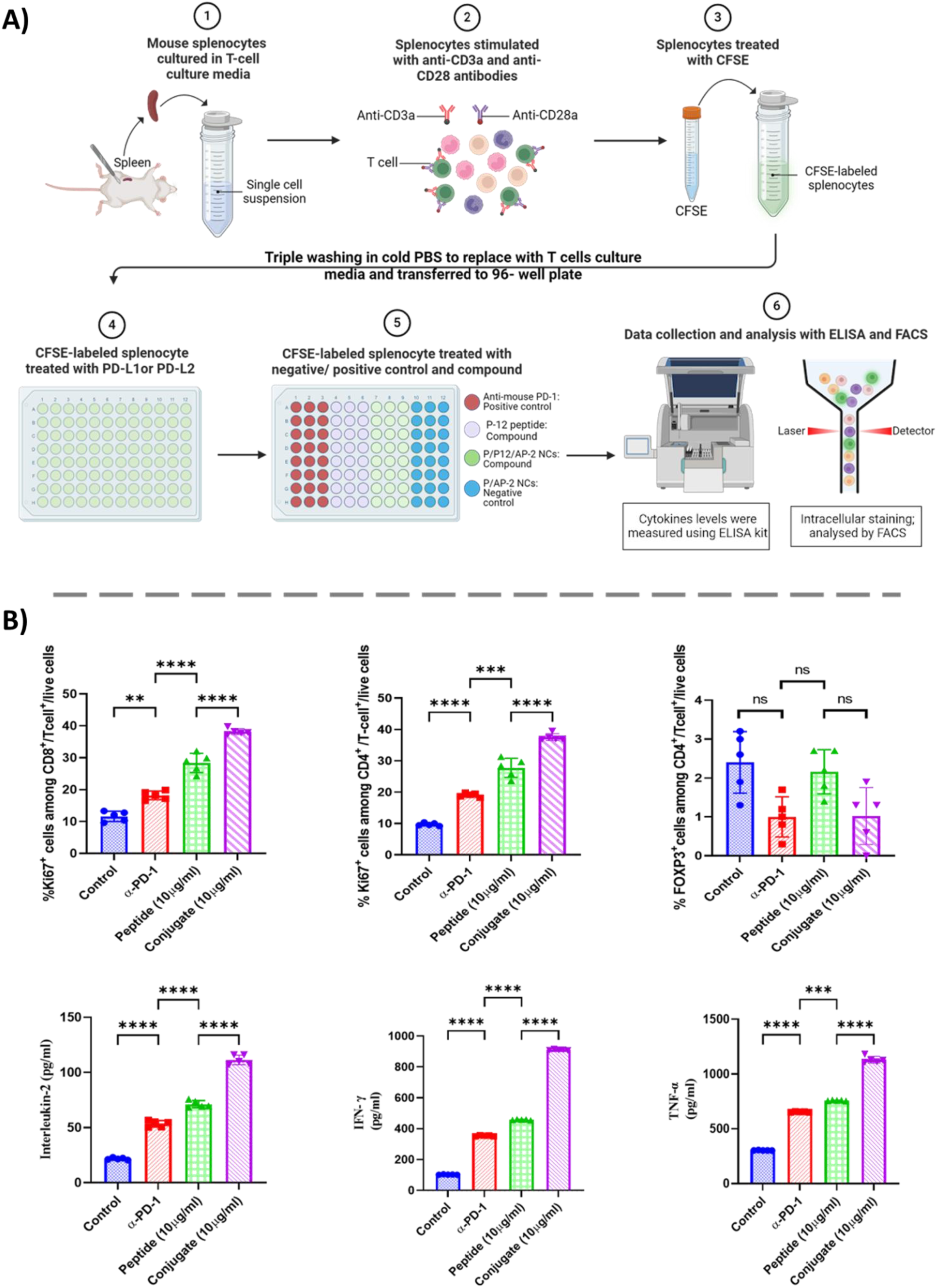
AP-2/P-12 nanoconjugates (NCs) modulate T lymphocyte proliferation and proinflammatory cytokine signaling. A) Schematic representation of the quantitative comparative T lymphocyte proliferation rescue analysis with Flow cytometry and ELISA. B) Subset-specific analysis of CD4+ and CD8+ T lymphocyte proliferation in response to anti-PD-1 antibody, P-12 peptide, and AP-2/P-12 NCs. Data are presented as mean ± standard error of the mean (SEM). Statistical significance was determined using one-way ANOVA followed by post-hoc tests, with ** p < 0.01, *** p < 0.001, and **** p < 0.0001 indicating significance levels between treatment groups. The results suggest a synergistic effect of the AP-2/P-12 NCs in promoting T cell activation and proliferation, which directly enhance anti-tumor immune responses.

### 2.4 AP-12/P-12 NCs induced T cell-mediated cytotoxicity of 3D GBM spheroids Co-culture

The Calcein-EtBr assay evaluated T-cell mediated cytotoxicity in 3D GBM spheroids. In this study, humanoid 3D GBM tumor spheroids were constructed using three distinct glioma cell lines (U87, LN 229, and PDM 140) and co-cultured with peripheral blood immune cells from healthy donors (HD-PBMCs), which were predominantly T-cells activated by anti-CD3/CD28 antibodies. The 3D GBM spheroids and T-cells were treated with saline, free P-12-pep, and AP-12/P-12 NCs to assess their cytotoxic effects. Treatment with AP-12/P-12 NCs resulted in a significant increase in red fluorescence intensity (indicating dead cells) in all three 3D GBM spheroids (U87, LN 229, and PDM 140), accompanied by an enhanced green fluorescence intensity (representing live cells) to red fluorescence intensity ratio compared to the free P-12-pep and control groups. This suggests that AP-12/P-12 NCs enhance T-cell cytotoxicity against 3D GBM spheroids (Fig. 5A).

**Figure 5.**
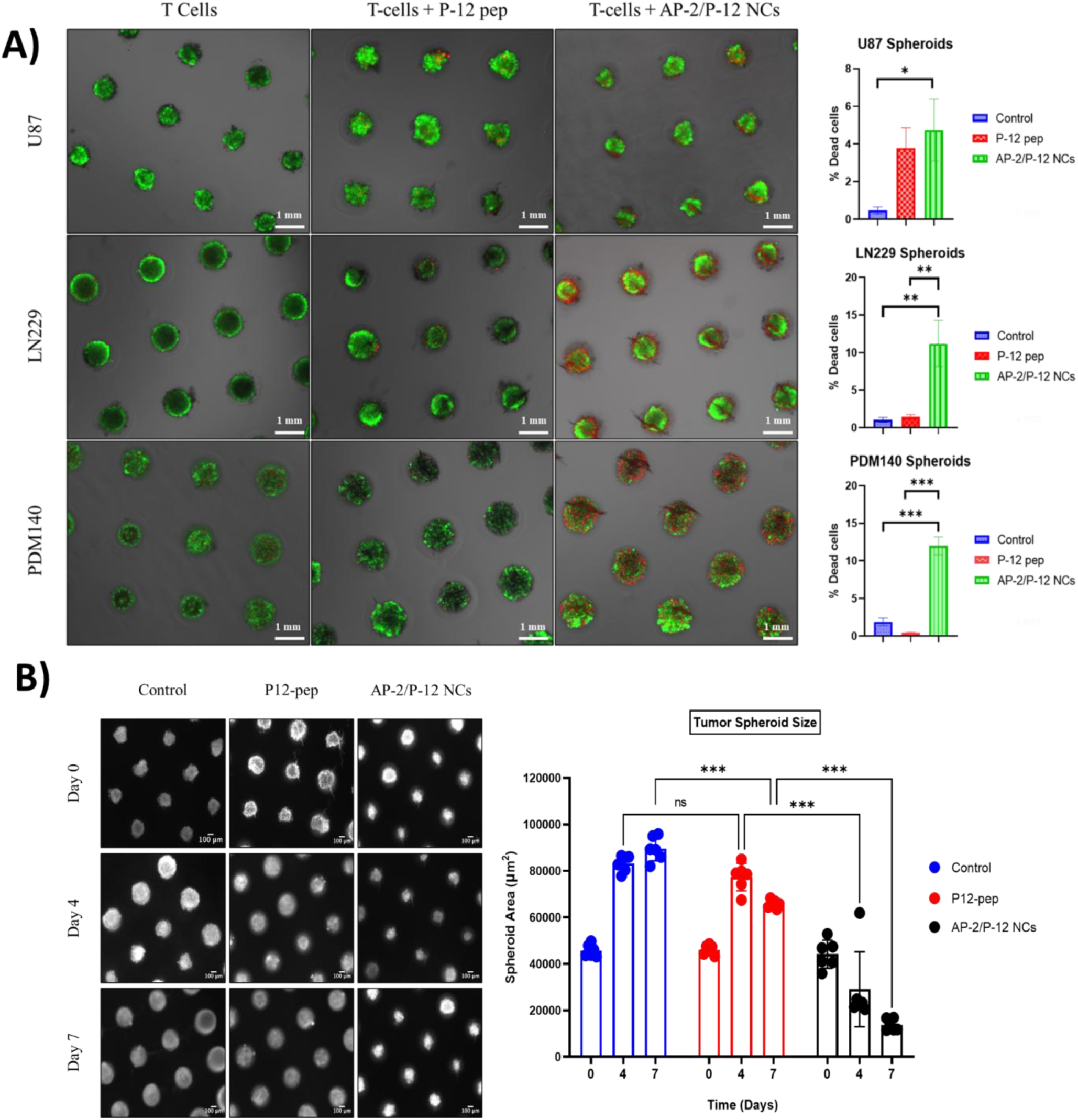
AP-2/P-12 nanoconjugates (NCs) enhance T cell-mediated cytotoxicity against glioblastoma multiforme (GBM) spheroids and inhibit their growth. A) Visualization of T cell-mediated cytotoxicity: Confocal microscopy images demonstrating the cytotoxic effects of T cells on GBM spheroids in the presence of P12 peptide and AP-2/P-12 NCs. Live cells are visualized using calcein AM (green fluorescence), while dead cells are identified by ethidium bromide (EtBr) staining (red fluorescence). This dual-fluorescence assay allows for the simultaneous quantification of live and dead cells within the spheroids. This provides a direct measure of T cell-mediated cytotoxicity. B) Temporal analysis of 3D tumor spheroid growth: Quantitative assessment of GBM spheroid size over 7 days when co-cultured with T cells in the presence of P12 peptide alone or AP-2/P12 NCs using brightfield microscopy. A reduced growth rate of spheroid size in the AP-2/P-12 NC treatment group over the 4 days can be observed. Data are presented as mean ± standard error of the mean (SEM). Statistical significance was determined using one-way ANOVA followed by post-hoc tests, with ** p < 0.01, *** p < 0.001, and **** p < 0.0001 indicating significance levels between treatment groups.

**Figure 6.**
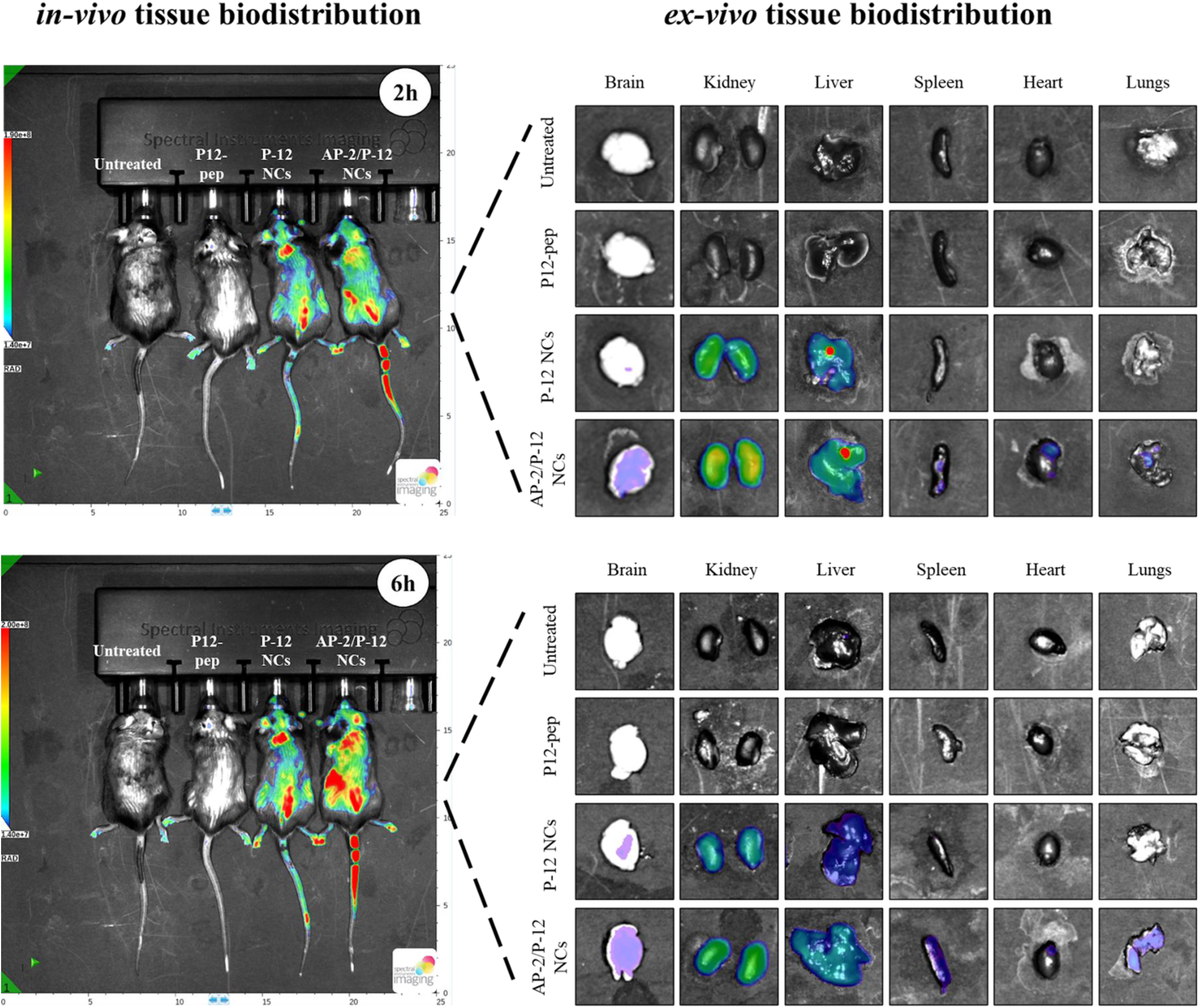
In vivo whole body and ex vivo tissue biodistribution at 2 h and 6 h after intravenous (i.v.) administration of AP2/P12/NCs. Biodistribution data at 2 and 6 hours post-i.v. Administration shows the sustained presence of AP-2/P-12 NCs in the brain.

### 2.5 AP-12/P-12 NCs generate T cell-mediated growth inhibition of 3D GBM spheroids

To evaluate the efficacy of AP-12/P-12 NCs inhibiting the growth of tumor spheroids, the volume of human-based U87 cells 3D spheroids was measured using bright-field images taken at various time points from day 0 to day 7. NCs containing the P12 exhibited a reduction in spheroid growth compared to the untreated control group. Interestingly, by day 7, spheroids treated with P12 peptide alone were larger than those treated with AP-12/P-12 NCs. Additionally, the administration of blank PBS alone did not significantly impact the growth of the tumor spheroids (Fig. 5B).

### 2.6 Biodistribution and histopathological toxicity studies of AP-12/P-12 NCs in C57BL/6 mice

The in vivo trafficking of nanoconjugates was determined using an In vivo imaging system (IVIS) in healthy C57BL/6 using FITC dye as a fluorescent probe. The mice were injected with saline, and free FITC-P-12 peptide showed no significant signal intensity at 2 and 6 h. On the contrary, upon administration of AP-12/FITC-P-12 NCs, an intense signal was observed throughout the body in 2 and 6 h. To better delineate the organ distribution of the nanoconjugates, the animals were euthanized at 2 and 6 h, and *ex vivo* signals of majorly excised organs (brain, kidney, liver, spleen, heart, and lungs) were determined. Meanwhile, saline and free P-12 peptide-injected animals showed no significant signal in any organ. Upon administration of AP-12/P-12 NCs, an intense signal was seen in the brain at 2 and 6 h compared to the P-12 NCs and free P-12-pep group. The signal was also observed in other organs, including the kidney, liver, spleen, and lungs, at both 2 and 6 h. AP-2/P-12 NCs penetrate into the brain more than P-12 nanoconjugate without AP-2. Both nanoconjugates also distribute into multiple viscera and promote longer drug bioavailability than free P-12 peptides. To determine the safety and toxicity profile of AP2/P12/NCs, we performed a toxicity study at 10 mg/Kg (low dose) and 20 mg/Kg (high dose). The H&E histopathological performed in major murine organs, including the brain, heart, and spleen, for any dose-dependent tissue toxicity effect of our AP-12/P-12 NCs with or without the conjugation of AP2. Mice were dosed with normal saline, 10 mg/kg, or 20 mg/kg of either variant of AP-12/P-12 NCs for 15 days and were then sacrificed. Overall, no adverse cellular adaptations were noted such as hyperplasia, fibrosis, or necrosis, with the administration of both concentrations compared to control (Supplementary Fig. S4).

## 3. Discussion

Glioblastoma multiforme (GBM) is a highly aggressive cancer characterized by a dauntingly limited range of therapeutic options. This limitation is primarily due to two factors: the TiME within the brain and its associated blood-brain barrier (BBB), which obstructs the penetration of approximately 90% of small-molecule drugs^19^, and the profoundly immunosuppressive tumor microenvironment (TiME) found within the brain^20^. Although immune checkpoint inhibitors such as anti-PD-L1, anti-PD-1, and anti-CTLA-4 have shown substantial therapeutic benefits in “hot” tumors—with higher lymphocyte infiltration—clinical trials for brain cancers often report suboptimal efficacy due to the "cold" nature of GBM, which is characterized by low immune cell infiltration at the tumor region^21^. To address these challenges, our research group has explored advanced drug delivery systems employing nanoconjugates (NCs) to facilitate receptor-mediated, peptide-functionalized nanoconjugates (NCs) therapy capable of circumventing these critical barriers. Our approach focused on developing brain-specific receptor-targeted peptides and antibody molecules using nanoconjugate technology to deliver immunotherapeutic agents across the BBB directly to the GBM. Dr. Holler’s research group previously synthesized PMLA polymer at Cedars-Sinai Medical Center which was used in this study. PMLA is advantageous due to its complete biodegradability to CO_2_ and H_2_O within cellular mitochondrial metabolic pathways, addressing the common biodegradability concerns of synthetic polyester polymers. For our study, we utilized 50 kDa-100 kDa β-PMLA polymers. One notable limitation of using purified β-PMLA lies in the complexity of chemical synthesis, which often results in α,β racemic mixtures and challenges in reproducibility^22^. This issue was successfully addressed by isolating and purifying a naturally derived PMLA polymer, enabling large-scale synthesis for further investigations^23^. Dr. Holler’s research findings demonstrated that PMLA was previously used for conjugation of several chemotherapeutics, oligonucleotides, and antibody-based nanoconjugates, with improved efficacy in animal models of brain cancer^14,24,25,29,30^. PMLA NCs possess intrinsic chemical properties that facilitate effective BBB penetration and targeted delivery within the tumor microenvironment, thus representing a compelling platform for developing immuno-nanotherapeutics. In previous work, it was demonstrated that combination therapies utilizing antibodies (anti-PD-1 and anti-CTLA-4) conjugated to PMLA yielded nanoimmunoconjugates (NICs) targeting GL-261 orthotopic mouse tumor models, resulting in reduced tumor volumes and significantly improved survival of glioma bearing animals ^23,31^. However, due to the immunotoxicity known for the therapeutic antibody in clinic, anti-PD-1 and anti-CTLA4 Abs were used with premedication to reduce the immunotoxicity after I.V. nano immunoconjugates administration^24^. Therefore, for greater efficacy with low toxicity, translational potential and cost considerations, here we have effectively designed an antibody-free, dual peptide-functionalized immuno-nanoconjugate incorporating a small-sized, cysteine-modified Angiopep-2 peptide (AP-2) selective for the LRP-1 receptor binding on BBB. We covalently linked this peptide to a PMLA-tri-leucine polymer nanoconjugate. Specifically, we replaced checkpoint-blocking antibodies with the highly effective AUNP-12 peptide (P-12) (KD; 0.41 nM), which targets the PD-L1 receptor. P-12 has already shown promising efficacy in preclinical models, including the B16F10 melanoma model, and was effective in rescuing T cells from PD-L1 and PD-L2 receptor interaction pathways. Our PMLA-based NCs incorporated cysteine-modified AP2 and P-12 peptides, successfully targeting both LRP-1 for transcytosis through BBB and PD-L1 within the GBM tumor-immune microenvironment. The AP2 peptide enhanced the NCs’ endocytosis via receptor-mediated mechanisms, improving cellular uptake in human brain endothelial cells and 3D-U87 cell spheroids. Notably, deeper analysis of the spheroid models revealed higher AP-2/P-12/RhB NCs’ endocytic uptake at the tumor periphery than at the necrotic core. Furthermore, our flow cytometry immune cells results suggest that the P12 peptide, when conjugated to our NCs, can inhibit immune checkpoints and reinvigorate T cell proliferation by blocking interaction with PDL1/PDL2 protein^5^. We observed enhanced immune cell activation within the GBM TiME compared to treatment groups lacking the nanoconjugate system. It should be noted that AP2/P12 NCs induced the highest proliferation of CD4+ and CD8+ T cells compared to other treatment modalities. At the same time, regulatory T-cell (T reg) expansion remained stable across all groups, suggesting that AP2/P12 NCs exhibit inherent immunogenic properties that stimulate immune cell activation without compensatory increases in regulatory populations. The ELISA analysis revealed significant elevations in pro-inflammatory cytokines, including IL-2, IFN-γ, and TNF-α, compared to controls. However, robust investigations into the contributions of the PMLA backbone and the LRP1-mediated delivery mechanism in modulating inflammatory signaling are warranted in subsequent studies. The AP2/P12 NCs effectively penetrated in an in vitro BBB model, demonstrating biodistribution in murine brains with minimal off-target effects. While promising, our current study presents directions for future research. Although our findings suggest that AP2/P12 NCs can provide significant immunodrug delivery across the BBB without noticeable toxicities, confirmation of efficacy necessitates further studies, especially with repeated dosing. Most importantly, we must evaluate the potential immunologic advantages of the PMLA backbone by including dedicated controls in future efficacy investigations to elucidate the distinct therapeutic potentials of our nano-delivery systems separate from drug or target mechanisms.

## 4. Conclusion

In conclusion, the AP2/P12 NCs emerge as a promising strategy to address GBM treatment’s formidable challenges. This research lays the ground for potential applications extending beyond GBM to encompass other difficult brain metastases and inflammatory brain disorders. Most importantly, our work reinforces the feasibility of developing scalable nanoconjugate production that can be tailored for specific targets and paves the way for innovative and personalized immunotherapeutic strategies. With further optimization, the AP2/P12 NCs show great promise as a transformative approach for treating GBM.

## 5. Materials

Poly β-L-malic acid (PMLA) with a molecular mass of ∼67,000 g/mol (as determined by SEC-HPLC using polystyrene sulfonate standards) was obtained and purified from the culture broth of *sp. Physarum polycephalum* M3CVII, as described previously^26^. The cysteine-modified Angiopep-2 peptide (AP-2) (sequence: H-TFFYGGSRGKRNNFKTEEYC-OH) was obtained from NovoPro Bioscience Inc. (Shanghai, China). Trileucine (LLL) was sourced from Bachem (CA, USA). Mal-PEG_3400_-Mal, mPEG_5000_-NH_2_, and Rhodamine Red™ C2 Maleimide (RhB) were purchased from Thermo Fisher Scientific (CA, USA). PD-10 columns were acquired from GE Healthcare (IL, USA). N,N’-Dicyclohexylcarbodiimide (DCC), N-Hydroxysuccinimide (NHS), trifluoroacetic acid (TFA), cysteamine (2-mercaptoethylamine, MEA), dithiothreitol (DTT), dimethylformamide (DMF), 3-(2-pyridyldithio)propionic acid (PDP), triethylamine (NEt3), and other chemicals were obtained from Sigma Aldrich (St. Louis, MO, USA).

## 6. Methods

### 6.1. Synthesis of Angiopep-2-PEG3400-maleimide (AP-2) pre-conjugate (1)

The Angiopep-2-PEG_3400_-maleimide (AP-2) pre-conjugate was prepared by dissolving Maleimide-PEG_3400_-Maleimide (3400 g/mol; 7.4 mg, 2.2 µmol, 1.05 equiv.) in 500 μL phosphate buffer (100 mM, pH 6.3) supplemented with 2 mM of EDTA. Subsequently, a solution of cysteine-modified Angiopep-2 peptide (H-TFFYGGSRGKRNNFKTEEYC-OH) (cys-AP-2) (5 mg in 500 µL of phosphate buffer; pH 6.3) was added to the maleimide mixture dropwise at 4 °C. The reaction was stirred for 1 h under ambient conditions, followed by purification using a PD-10 column (MWCO. 3KDa) and freeze-dried. The product was reconstituted in phosphate buffer (10 mg/mL, pH 6.3) to synthesize PMLA conjugates.

### 6.2 Synthesis of P-12 peptide nanoconjugates (P-12/NCs, FITC-P-12/NCs) conjugate

The P-12 NCs were synthesized using carbodiimide coupling. Briefly, 320 mg of PMLA, which was previously synthesized ^15, 23–29^ and dissolved in 3 mL of DMF. Subsequently, a mixture of 320 mg of N-hydroxysuccinimide and 592 mg of N, N’-dicyclohexylcarbodiimide (DCC) in 3 mL of dimethylformamide (DMF) was added dropwise to activate the PMLA, followed by stirring for 2 h at room temperature. Afterward, a separate DMF solution of tri-leucine (394.4 mg), P-12 peptide (2.15 mg) or FITC-P-12 along with trifluoroacetic acid (101 μL) was added with an interval of 1 h. After 2 h, triethylamine (NEt3) was added to maintain the succinimide integrity. Thereafter, the reaction mixture was stirred for 12 h, followed by adding 31.36 mg of mercaptoethylamine hydrochloride (MEA.HCl) and NEt3 (1:1 molar ratio of MEA). The reaction mixture was allowed to stir for 3 h to yield thiol pendant polymer PMLA/LLL/P12 or FITC-P12/MEA. Afterward, the unreacted thiol (SH) pendant groups were protected with DL-dithiothreitol (DTT) (56 mg) to yield the final polymer. The polymers were purified using a PD-10 column and subsequently lyophilized. The product was stored at −80 °C until further use.

### 6.3. Synthesis of AP-2/P-12 NCs or AP-2/P-12/RhB NCs

The AP-2/P-12 NCs were synthesized by dissolving 30 mg of P-12 NCs or FITC-P-12 NCs (130 g/mol, 9.19 µmol) in 3 mL of degassed PBS (150 mM, pH 6.3) in a glass vial. Thereafter, AP-2-PEG_3400_-maleimide (14.24 mg, 2.37 µmol) was dissolved in 1.3 mL of PBS (150 mM, pH 6.3) and added dropwise to the reaction mixture and allowed to stir for 3 h under ice-cold (4 °C) conditions. Thereafter, Rhodamine Red™ C2 Maleimide (0.1 mg, 680.79 g/mol, 0.146 µmol) was added in dark conditions and stirred for 3 h at 4 °C. Subsequently, 120 µL of PDP (10 mg/mL solution in DMF) was introduced to cap the free thiol groups and the solution was stirred for an additional hour. The crude reaction mixture was then purified using a PD-10 column and PBS as a solvent. The amount of rhodamine-conjugated to the PMLA backbone was analyzed using spectrofluorimetric and HPLC analysis, followed by freeze-drying. The samples were stored at - 80 °C until further use.

### 6.4. Polymer and Peptide Quantification

The synthesis of nanoconjugates was monitored through size-exclusion chromatography (SEC-HPLC) using the Agilent instrument with a Hitachi L-2455 detector, and data was analyzed using EZChrome Software. The samples were eluted on a Polysep 4000 SEC column at a 1 mL/min flow using PBS (pH 7.4, 150 mM) as a mobile phase. The molecular weight of PMLA and nanoconjugates were determined using polystyrene sulfonate as standards detected at λ_max_ 220 nm (Fig. 2C). Reverse-phase HPLC was used to quantify the amount of unconjugated P-12 peptide within the reaction mixture (using an indirect method) for the determination of the number of P-12 peptide units attached to PMLA polymeric backbone.

### 6.5. Particle size (PS), polydispersity index (PDI), and Zeta Potential (ZP, ζ) Characterization

The nanoconjugates were prepared at a concentration of 2 mg/mL (PBS, pH 7.2), and their particle size (PSA; nm), polydispersity Index (PDI), and zeta potential (ZP; **ζ**) were determined using a Malvern Nano-ZS zeta sizer (Malvern Panalytical, UK).

### 6.6. 3D Tumor Spheroid Formation

With slight modifications, the humanized 3D-brain tumor spheroids were prepared using 3D Petri Dishes® technology from MicroTissues Inc (Sigma-Aldrich, MO, USA). Briefly, 2% sterile agarose in 0.9% NaCl was heated at ∼70 °C and poured to a 12-series micro mold and allowed to cool down under ambient conditions and were made sure for the complete removal of bubbles from the mold. The resulting 12-series agarose micro-molds containing agarose gel were transferred to 12-well plates and equilibrated with DMEM containing 10% FBS and 0.1% Penicillin-Streptomycin mixture. This equilibration was repeated twice. The trypsinized cell (U87MG, PDM140 and LN229 cells) suspension was seeded into the micro-molds at a density of ∼50 × 10^3^ cells per well. The cells were allowed to settle down for ∼10 min, followed by adding 2.5 mL media to each well and kept in an incubator for their growth. The spheroid formation within the 3D Petri Dish® was tracked daily using an inverted microscope until further experimentation.

### 6.7. PBMCs Isolation and T Cell Isolation, Activation, and Culturing

Human PBMCs were isolated from healthy donors’ blood by density gradient centrifugation using the Ficoll-Paque PLUS method. Briefly, the blood was diluted with phosphate-buffered saline (PBS) (1:1 v/v), layered on Ficoll-Paque media, and centrifuged at 400 × g for 30 min. To remove RBC, the PBMC with RBC lysis buffer was added and incubated for 2 min at 4°C, washed twice and re-suspended with PBS. Isolated PMBCs were immediately enriched by Phorbol 12-myristate 13-acetate (PMA) (5ng/mL) for 24 h. For activation of PBMCs, isolated cells were cultured in 6 well-plates in RPMI-1640 medium with 10% fetal bovine serum (FBS), 100 IU/mL penicillin, and 100 µg/mL streptomycin. Thereafter, the PBMCs were incubated with human-specific anti-CD3a and anti-CD28 antibodies (1 μg/mL each) for 72 h at 37 °C with 5% CO_2_ supplemented with 50 IU/mL of recombinant human IL-2 (Novus Biologicals, Littleton, CO, USA). The activated PBMCs were utilized for further experiments.

### 6.8. Spheroid Live/Dead assay

Spheroid Live/dead assay was performed using humanized 3D-brain tumor spheroids (U87MG or LN229 or PDM140) was prepared as described in section 4.4. The spheroid was co-cultured with CD-3 and CD-28 expressing PBMCs and treated simultaneously with media, free P-12 pep (P-12-pep), and AP-2/P-12 NCs for 7 days. After that, the supernatant was removed and tumor spheroids were washed gently with Ice-cold PBS twice. Thereafter, the spheroids were stained using Calcein AM/ ethidium bromide Live/Dead staining kit (Invitrogen, cat. # L3224) to stain live cells with green fluorescence and ethidium homodimer-1 to label dead cells with red fluorescence, as per manufacturer protocol. Briefly, the Calcein AM (2 μM) and Ethidium Homodimer (4 μM) staining solution was added to the spheroids and incubated for 10 min at 37 °C under dark conditions, then washed with sterile PBS twice. The representative visualization was achieved thereafter using a fluorescent microscope with green and red fluorescence filters. The recorded data was quantified and assessed using the Fiji plug-in within the ImageJ software.

### 6.9. In vitro Cellular uptake of Nanoconjugates in BBB-transwell-3D spheroids system

Immortalized human cerebral microvascular endothelial cells (HBMEC) were cultured in Transwell plates to create an *in vitro* blood-brain barrier (BBB) model, as described earlier^27^. The monolayered human brain endothelial cells (HBMEC) cells were cultured in differentiation medium (EBM-2 supplemented with 5% FBS, 1% penicillin/streptomycin, 1.4 µM hydrocortisone, 5 µg/mL ascorbic acid, 1% lipid concentrate, 10 mM HEPES buffer, and 1 ng/mL bFGF) for 3 days at 37 °C with 5% CO2 and saturated humidity. Subsequently, the medium was switched to the growth medium (EBM-2 supplemented with 5% FBS, 1% penicillin/streptomycin, 1.4 µM hydrocortisone, 5 µg/mL ascorbic acid, 1% lipid concentrate, 10 mM HEPES buffer, and 10 mM LiCl) for 6-7 days, with medium changes occurring every 3^rd^ day. 50 × 10^3^ cells were seeded on the apical side of 12-well polystyrene transwell plates with 0.4 µm pores and 1.12 cm^2^ inserts (Corning™ 3402, Cat. 07-200-157). In vitro, a blood-brain barrier (BBB) model was created by culturing human brain endothelial cells (HBMEC) 50 x10^3 cells on the top of the fibronectin (2 μg/cm^2^) layer as an extracellular matrix for 7 days. The in-vitro BBB model was validated through the characteristic BBB-tight junction proteins including ZO-1, Occludins, claudins, and VEGF using a fluorescent microscope (Supplementary Fig. S2). Thereafter, the penetration capability of AP-2/P-12 NCs was determined through a transwell BBB-based 3D spheroid organ on-chip model (in vitro BBB and 3-D tumor Spheroids) to better mimic the in vivo BBB and tumor environment. After that, the media, free Rhodamine (RhB), AP-2/P-12/RhB NCs were added to the apical compartment at a concentration equivalent to 46.7 ng/mL for 4 h. Post-treatment, the media was discarded and the BBB (in apical compartment) and spheroids (in basal compartment) were washed gently with PBS twice, followed by staining with Hoechst and cell tracker (for spheroids only) under dark conditions. The representative images (Hoechst, cell tracker, rhodamine, and overlay) were taken using fluorescent confocal microscopy, and the data was interpreted using the Fiji plug-in embedded within ImageJ software.

### 6.10. Efficacy of nanoconjugates on T cell proliferation inhibiting PDL1

T-cell proliferation efficiency of AP-2/P-12 NCs was determined by inhibiting PD-1/PD-L1/L2 complex inhibition. Wherein the proliferation of T-cells expressing CD4, CD8, and inflammatory cytokines. Mouse splenocytes were isolated to single-cell suspension, as described elsewhere^12^. The splenocytes were cultured in cell culture media (RPMI with 10% FBS + 2mM L-glutamine + 0.5 mM of 2-mercaptoethanol + 2 mM Nonessential Amino acid solution + 10,000 IU/mL PEN-Strep). The splenocytes were treated with specific anti-CD3a and anti-CD28 antibodies (1 μg/mL each) and incubated for 72 h at 37 °C with 5% CO2. After 72 h, the splenocytes (10 × 10^6 cells/mL) were treated with 1 mmol/L carboxyfluorescein succinimidyl ester (CFSE) for 30 min at 37 °C in 1 mL 0.1% BSA in PBS. After this, CFSE was removed and cells were washed with PBS. The process was repeated 4 times and incubated on ice for 5 min to remove any remaining free CFSE dye. After that, CFSE-labeled splenocytes were seeded in a 96-well plate at a density of 1 x 10^5 cells/well and exposed to a solution containing recombinant PD-L1 and PD-L2 (10 nmol/L each) protein, simultaneously treated with fresh media (normal control), 100 nmol/L InVivoMAb anti-mouse PD-1 (CD279) (positive control) (α-PD-1), 10 µg/mL of free P-12 peptide (P-12-pep), AP-2/P-12 NCs (equivalent to 10 µg/mL of P-12 peptide). The cells were incubated for 72 h and centrifuged at 2000 rpm at 4°C for 5 min. The media was collected and cytokines, including IL-2, TNF-α, and IFN-γ cytokines levels were measured at 450 nm using an ELISA assay kit. Simultaneously, the cells were harvested and washed thrice with PBS and resuspended in a cold FACS buffer. The cells were incubated with fluorescently labeled antibodies targeting CD8+, CD4+, and T-reg cells for 30 min at 4°C in dark conditions. Thereafter, the cells were washed thrice with FACS buffer, and fixation/permeabilization buffer was added to enable intracellular staining with anti-Ki67 and FoxP3 antibodies. The labeled cells were washed twice and resuspended in FACS buffer. The analysis was carried out using FACS Calibur (FACS AriaIII), with a minimum of 50,000 total events were recorded. The cells positive for each marker were identified and the percentage of Ki67-positive cells was determined for interpretation using FlowJo software.

### 6.11. In-vivo and ex-vivo Biodistribution

Biodistribution studies of AP-2/P-12 NCs were performed in healthy C57BL/6 animals with 6-8 weeks of age using IVIS Imaging system, wherein the animals were treated with PBS, P-12-pep, FITC-conjugated P-12 NCs, FITC-conjugated AP-2/P-12 NCs at a dose equivalent to 10 mg/kg i.v. The animals were imaged at 2 h and 6 h intervals for in vivo imaging. Simultaneously, the animals were euthanized at 2 h and 6 h and their major organs (including brain, heart, lungs, liver, kidney, and spleen) were isolated for ex vivo tissue biodistribution analysis. The organs were promptly rinsed in ice-cold normal saline to remove blood, pat dried and *ex-vivo* organ imaging was performed using the IVIS Imaging system. This allowed us to determine the fate of nanoconjugates via in vivo administration among the different organs over time.

### 6.12. Animal safety and toxicity study

In this study, animals were divided into two dose levels of approximately 10 mg/kg and 20 mg/kg equivalent amount of P12 peptides, administered with nanoconjugates composed of different formulations: P-12 NCs and AP-2/P-12 NCs with a total of 21 animals. Each animal received a single intravenous dose. Mice were randomly assigned to receive either the nanoconjugates groups or a control group intravenously. Body weights were monitored twice weekly, and clinical observations were conducted at least twice daily during dosing and once daily thereafter. Mortality checks were performed twice daily lasting at least 6 h between each check. On day 15, blood samples were obtained via transcardiac aspiration for hematological evaluation, and vital organs (brain, heart, kidney, lung, liver, and spleen) were collected for histopathological analysis immediately after euthanasia.

## 7. Statistical Analysis

Data were analyzed using GraphPad software. Results are expressed as mean +/- s.e.m. or s.d. Data were analyzed by one-way ANOVA followed by post-hoc tests, with ** p < 0.01, *** p < 0.001, and **** p < 0.0001 indicating significance levels between treatment groups.

**Table 1.**
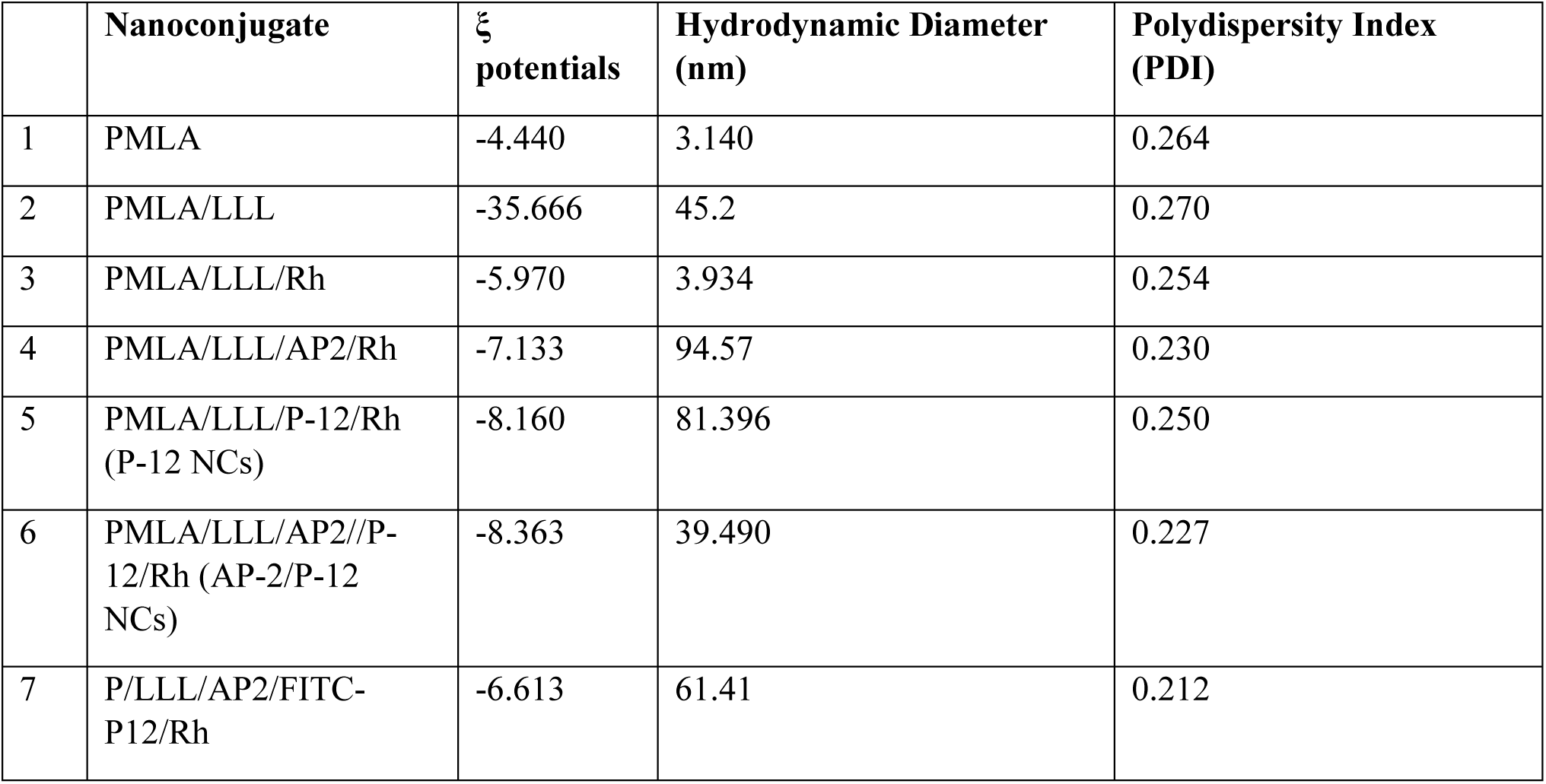
Hydrodynamic diameter, polydispersity index (PDI) and ξ potentials.

|  | **Nanoconjugate** | **$\xi$**<br>**potentials** | **Hydrodynamic Diameter**<br>**(nm)** | **Polydispersity Index**<br>**(PDI)** |
| --- | --- | --- | --- | --- |
| 1 | PMLA | -4.440 | 3.140 | 0.264 |
| 2 | PMLA/LLL | -35.666 | 45.2 | 0.270 |
| 3 | PMLA/LLL/Rh | -5.970 | 3.934 | 0.254 |
| 4 | PMLA/LLL/AP2/Rh | -7.133 | 94.57 | 0.230 |
| 5 | PMLA/LLL/P-12/Rh<br>(P-12 NCs) | -8.160 | 81.396 | 0.250 |
| 6 | PMLA/LLL/AP2//P-12/Rh (AP-2/P-12<br>NCs) | -8.363 | 39.490 | 0.227 |
| 7 | P/LLL/AP2/FITC-P12/Rh | -6.613 | 61.41 | 0.212 |

## 8. Acknowledgement

This work was supported by NIH R01 CA209921 (EH) grant “Glial tumor image-guided surgery and treatment” at year 5. Dr. Holler is PI on this grant.

