## Supplemental Files for "Antibody-Free Immunopeptide Nano-Conjugates for Brain-Targeted Drug Delivery in Glioblastoma Multiforme"

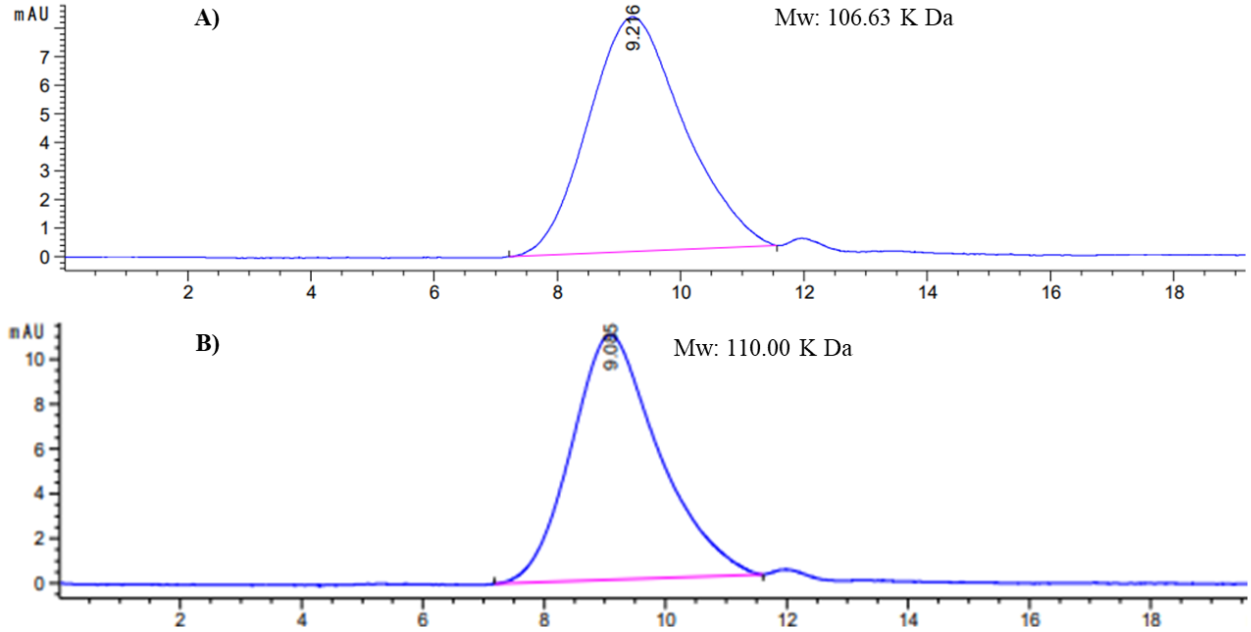


Figure S1. Characterization by using SEC-HPLC for determining the total molecular weight distribution of the synthesized A) PMLA/LLL/MEA/RhB NCs and B) PMLA/LLL/MEA/P12/RhB

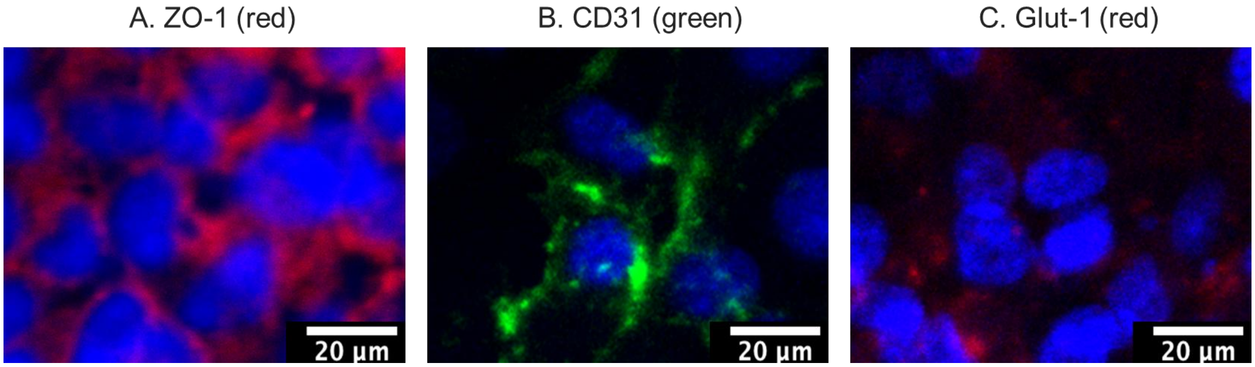


Figure S2. Confocal laser scanning microscopy (CLSM) analysis of HBMECs (seeded as a monolayer on fibronectin-coated trans wells) mimicking the BBB: A) ZO-1 (red) is a protein located on a cytoplasmic membrane surface of intercellular tight junctions, B) CD31 (green) (platelet endothelial cell adhesion molecule-1, PECAM-1), C) GLUT-1 (red) Facilitative glucose transporter. Scale bars represent 20 µm

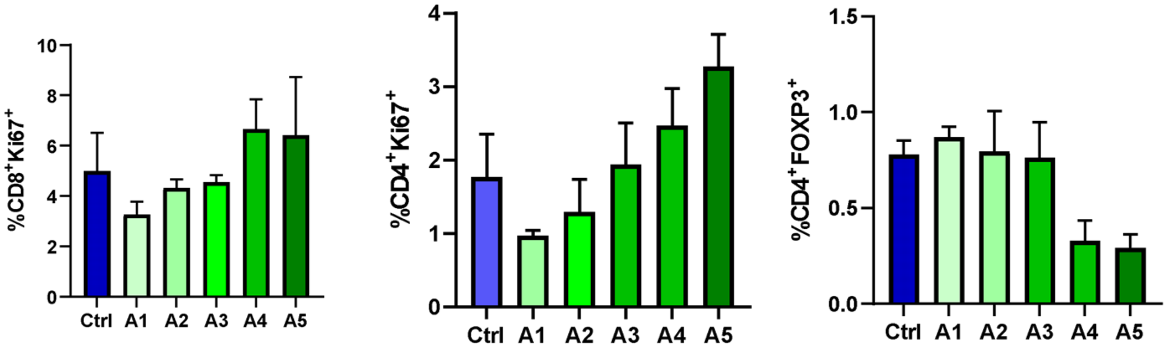


Figure S3: T cells proliferation with PDL1/PDL2 recombinant protein at various concentrations of P12 peptide from A1-A5 (nM/L): 1, 5, 10, 50, 100 respectively.

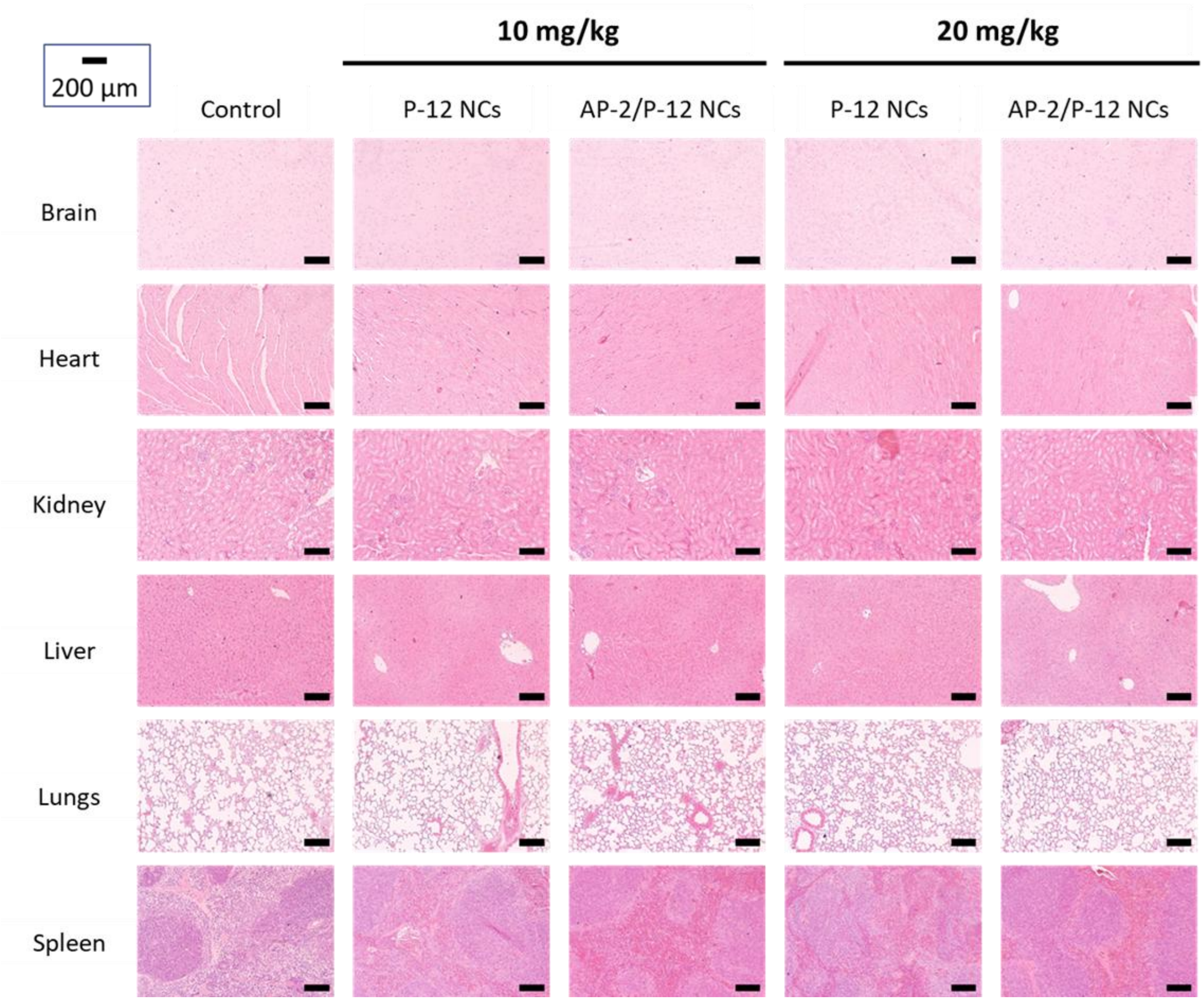

Figure S4: Histopathological assessment of organ toxicity using hematoxylin and eosin (H&E) staining. Results indicate that all treatment groups, including those receiving AP-2/P-12 NCs, exhibited tissue morphology comparable to untreated controls. No signs of acute or chronic toxicity, such as necrosis, fibrosis, or significant inflammatory responses, were observed in any examined organs. This suggests that AP-2/P-12 NCs demonstrate a favorable safety profile with no detectable organ toxicity.
